# TGFβ1-induced cell motility is mediated through Cten in colorectal cancer

**DOI:** 10.1101/339341

**Authors:** Abdulaziz Asiri, Teresa Pereira Raposo, Abdulaziz Alfahed, Mohammad Ilyas

## Abstract

Cten is a tensin which promotes epithelial-mesenchymal transition (EMT) and cell motility. The precise mechanisms regulating Cten are unknown, although Cten could be regulated by several cytokines and growth factors. Since Transforming growth factor beta 1 (TGF-β1) regulates integrin function and promotes EMT / cell motility, we investigated whether this happens through Cten signalling in colorectal cancer (CRC).

TGF-β1 signalling was modulated by either stimulation or knockdown in the CRC cell lines SW620 and HCT116. The effect of this modulation on expression of Cten, EMT markers and cellular function was tested. Cten role as a direct mediator of TGF-β1 signalling was investigated in a CRC cell line with a deleted Cten gene (SW620^ΔCten^).

When TGF-β1 was stimulated or inhibited, this resulted in, respectively, upregulation and downregulation of Cten expression and EMT markers. Cell migration and invasion were significantly increased following TGF-β1 stimulation and lost by TGF-β1 knockdown. TGF-β1 stimulation in SW620^ΔCten^ resulted in selective loss of the effect of TGF-β1 signalling on EMT and cell motility whilst the stimulatory effect on cell proliferation was retained.

These data suggested Cten may play an essential role in mediating TGF-β1-induced EMT and cell motility and may play a role in metastasis in CRC.

## INTRODUCTION

C-terminal tensin-like (Cten, also known as tensin4) is the member of the tensin gene family which comprises four members (tensin1, tensin2, tensin3, and Cten/tensin4). This protein family localises to the cytoplasmic tails of integrins at focal adhesion sites. Cten shares high sequence homology to the C-terminus of the other tensins with a common Src homology 2 (SH2) domain and phosphotyrosine binding (PTB) domain. Unlike the other tensin protein members (tensins 1-3), Cten lacks the actin binding domain (ABD) which results in an inability to bind to the actin cytoskeleton and is thought to play a critical role in cellular processes such as cell motility (1).

Cten is a putative biomarker in many cancers, acting as oncogene in most tumour types including the colon, breast, pancreas and melanoma, and it is particularly associated with metastatic disease (2). Cten expression is possibly upregulated through the activation of upstream signalling pathways since so far, no mutations or amplification of Cten in cancers have been documented. A study by Katz et al. showed that stimulation with EGF led to upregulated Cten expression at a post-transcriptional level only in breast cell lines, whereas others have shown that Cten is upregulated by the EGFR at both the transcriptional and post-transcriptional level (3, 4). Further reports suggested that Cten is regulated by KRAS in both CRC and pancreatic cancer cells (5). Cten expression was also shown to be negatively regulated by STAT3 in CRC cell lines, whereas others have found that Cten is upregulated by STAT3 in human lung cancer cells (6, 7). How Cten is activated and regulated in these tumours is unclear, nonetheless, multiple pathways seem to be involved, and it appears to be largely dependent on tissue type or context.

Transforming growth factor beta 1 (TGF-β1) is a polypeptide member of the growth factor family that plays a physiological role in the regulation of wound healing, angiogenesis, differentiation, and proliferation. TGF-β 1 can function as a tumour suppressor in normal epithelial cells and in the early stage of cancer. However, the growth inhibitory function of TGF-β1 is selectively lost in late stage cancer which results in an induction of cell migration, invasion and metastasis (8, 9). Previous studies have shown that TGF-β1 is involved in the regulation of EMT processes through numerous downstream pathways, including Ras/MAPK (10), RhoA (11), and Jagged 1/Notch (12). TGFβ1 has also been found to signal through FAK to upregulate EMT related mesenchymal and invasiveness markers and delocalise E-cadherin membrane (13). TGF-β1 has been shown to regulate several integrins including αV, β1, and β3 in glioblastoma, fibroblast, and kidney epithelial cells (14). Others have suggested that the positive regulation of integrin of αV, α6, β1, and β4 by TGF-β1 signalling is probably mediated via the activation of the TGF-β1/TGF-βRI/Smad2 signalling pathway (15). Furthermore, the TGF-β1 mediated Smad signalling pathway has been shown to play an important role in EMT associated with metastatic progression (10).

There are therefore several cellular functions and processed which are similarly regulated by Cten and TGF-β1. Both molecules seem to use FAK as a downstream messenger. However, a possible role of Cten in TGF-β1 mediated EMT and cell motility in CRC cells has not yet been postulated. Therefore, it was hypothesised that TGF-β may induce cell motility and promote EMT processes through the Cten signalling pathway.

## MATERIALS AND METHODS

### Cell Culture

This work was performed in CRC cell lines HCT116 and SW620, which were a kind gift from Prof Ian Tomlinson. This work was also carried out in SW620^ΔCten^ cell line which was previously established by our group (16). All cells were cultured in Dulbecco’s Modified Eagle’s Medium (DMEM) (GlutaMAX™ supplement, Thermo Fisher Scientific, Carlsbad, CA) antibiotic free supplemented with 10% foetal bovine serum (FBS) (Sigma) and maintained at 37°C in a 5% CO2 atmosphere. Cell line identities were authenticated by STR profiling.

### Cell Transfection

Cells were transfected with small interfering RNA (siRNA) targeted to TGF-β1 (CCA CCU GCA AGA CUA UCG ACA UGG A) or luciferase (CGU ACG CGG AAU ACU UCG A) as a negative control using Lipofectamine 2000 (Invitrogen) according to the manufacturer’s protocol. Cells were grown to 40-50% confluency and siRNA duplexes were added at a final concentration of 10 nM and 10 μl of Lipofectamine 2000. The cells were incubated with the transfection reagents for 6 hours and experiments performed 48 hours post transfection.

### Cell Treatment

In order to stimulate TGFβ1 signalling, cells were seeded in a six well plate and starved in serum free DMEM for 24 hours at 37°C prior to stimulation. Cells at 50-60% confluency were treated with 0–20 ng/ml Recombinant Human TGFβ1 (R&D Systems) in DMEM growth media (supplemented with 10% FBS), with a total volume of 2 ml per well. Cells were harvested after incubation for 48 hours.

### Western Blot

Cell lysates were prepared using RIPA lysis buffer (Thermo Fisher Scientific) supplemented with phosphatase and protease inhibitor (Thermo Fisher Scientific). Fifty micrograms of protein was added to NUPAGE LDS Sample Buffer (Thermo Fisher Scientific) containing 5% β-mercaptoethanol. The protein samples were heated to 90°C on a heat block for 5 minutes and cooled on ice for another 5 minutes. Following this, the protein samples were fractionated on a pre-cast 4– 12% NUPAGE Bis-Tris-HCl buffered (pH 6.4) polyacrylamide gel (Thermo Fisher Scientific) using the NUPAGE gel electrophoresis system with NUPAGE MOPS SDS Running Buffer (Thermo Fisher Scientific) at 125 V for 90 minutes. Proteins were then transferred onto PVDF membrane (GE Life Sciences) using the Trans Blot semi-dry transfer system (Biorad). Following blocking in 5% milk or 5% BSA in 0.1% tween PBS or 0.1% tween TBS (dependent on antibody diluents), membranes were incubated with optimally diluted primary antibodies overnight at 4°C (supplementary Table 1). Following washing, membranes were incubated with the appropriate anti-mouse or anti-rabbit secondary antibody for 1 h at room temperature (supplementary Table 1). The ECL prime detection kit (GE Life Sciences) was used for protein band visualisation using the C-DiGiT Blot Scanner (LI-COR, Lincoln, NE). Densitometric analysis of the bands was performed using ImageJ software. Pixel counts for each protein of interest were normalized to β-actin.

### Nuclear and Cytoplasmic Extraction

The nuclear and cytoplasmic fractions were extracted using the NE-PER Nuclear and Cytoplasmic Extraction Kit (Thermo Scientific) according to the manufacturer’s protocol. Briefly, cells were harvested by trypsinisation 48 hours post transfection and pelleted by centrifuging at 500 x g for 5 minutes. The pellet was then lysed in ice-cold CER I supplemented with protease inhibitor for 10 minutes, followed by addition of ice-cold CER II and incubation on ice for 1 minute. Cell lysates were then centrifuged at 16,000 x g for 5 minutes at 4°C and the supernatant containing cytoplasmic proteins was removed and stored for use. The insoluble pellet was resuspended in ice-cold NER supplemented with protease inhibitor and incubated on ice for 40 minutes and then centrifuged at 16,000 x g for 5 minutes at 4°C. Next, the supernatant containing nuclear fractions was transferred to a clean, pre-chilled tube. Thereafter, western blotting was performed as previously described.

### Immunofluorescence

Cells were seeded in multi chamber slides and incubated at 37°C for 24 hours post transfection or treatment. Cells were washed in PBS for 5 minutes on a shaker and fixed with 1:1 acetone/methanol for 20 minutes at −20°C. Following this, cells were washed three times in PBS for 5 minutes each and blocked with 1% BSA and 22.52 mg/mL glycine in PBS-T (PBS+0.1% Tween 20) at room temperature for 60 minutes. After blocking, the primary antibody diluted in 1% BSA+PBS-T was incubated with cells overnight with gentle rotation at 4°C (supplementary Table 2). Then, cells were washed three times in PBS for 5 minutes each and incubated with Alexa Fluor 488 (green) or Alexa Fluor 568 (17) secondary antibody (Thermo Fisher Scientific) diluted in 1% BSA+PBS-T with gentle rotation at room temperature in the dark for 60 minutes. Cells were then washed three times in PBS-T for 5 minutes in the dark and incubated with 1x DAPI (1:5000) (Sigma) in PBS with gentle rotation for 30 minutes. Following this, cells were again washed a further 2 times in PBS for 5 minutes in the dark and mounted with a drop of fluorescein mounting media (Sigma) on glass slides. The cells were viewed using the Leica Microsystems confocal microscope at x40 objective and Leica Application Suite X software was used for image acquisition.

### Cell Viability Assay

PrestoBlue Cell Viability Reagent (Thermo Fisher Scientific) was used as an indirect method to measure the total number of live cells. Briefly, 5 x 10^3^ cells were seeded in a 96 well plate and allowed to attach for 24 hours. Following this, 100 μl of PrestoBlue Cell Viability Reagent was added to the cells and without cells as control, and incubated for 1 hour at 37°C. Following incubation, the fluorescent unit (OD) of each well of the 96 well plate was then measured using the BMG FLUOstar Optima Plate reader (540 nm/ 590 nm). Further readings were taken at 48 and 72-hour time points. The blank fluorescence reading was subtracted from each experimental fluorescence reading and the blank corrected values were then normalised to the 24 hour time point.

### Transwell Migration and Invasion Assays

The changes in cell migration were assessed in 24 well plates using the Transwell system (Corning, Corning, NY). Cells were treated with mitomycin C (10 μg/ml) for 3 hours and washed 3 times with PBS to inhibit cell proliferation. The Transwell inserts (6.5 mm diameter; 8 μm pore size) were incubated in DMEM at 37°C for 1 hour prior to use. Following this, 600 μl of DMEM (20% FBS) was added to the outer wells of the Transwell plate and the Transwell inserts placed inside. A total of 1 × 10^5^ cells in DMEM (10% FBS) were seeded onto the Transwell insert. The plate was incubated at 37°C for 24 hours. Following this, the cells that had migrated through to the bottom of the outside well, using the higher FBS concentration as a chemoattractant, were manually counted. Triplicate wells were seeded for each experimental condition. The Transwell invasion assay was performed according to this protocol with the exception that 2 × 10^5^ cells were seeded onto a Transwell insert coated in matrigel reduced growth factor (0.3 mg/ml, Corning) and cells allowed to migrate for 48 hours prior to counting.

### Wound Healing

Wound healing” scratch assay was performed as an alternative assay to assess cell migration. Briefly, 5-7 x 10^5^ cells /ml were seeded into culture-insert 2 well (Ibidi) attached in 6 well plates and incubated for 24 hours at 37°C until they reached confluency. After cell attachment, the culture inserts were removed, and the cells were treated with mitomycin C (10 μg/ml) for 3 hours to inhibit proliferation. Cells were cultured under normal conditions, and the pictures were taken at time points 0, 24, or 48 using an inverted microscope (Nikon) at 10x magnification. The width of the cell free gap was approximately 500 microns (+/− 50 microns) at time 0 hours. Experiments were performed in duplicate and on at least on two separate occasions. Cell migration was assessed by measuring the remaining open area of the wound by ImageJ software.

### Statistical Analysis

Results were tested for a normal distribution, and the unpaired t-test or analysis of variance (ANOVA) statistical tests were applied using GraphPad Prism (version 6).

## RESULTS

### TGF-β1 Regulates Cten Expression

Both TGF-β1 and Cten induce EMT and cell motility and are associated with integrin signalling. Thus it was of interest to determine whether Cten expression is under the regulation of the TGF-β1 signalling pathway in CRC cells. To investigate this, SW620 cells were treated with different concentrations of TGF-β1-human recombinant protein (R&D systems) from 0 to 20 ng/ml for 48 hours and the changes in protein level of Cten and its downstream targets, ROCK1, N-cadherin, E-cadherin, Src, and Snail were evaluated by western blot. SW620 cells showed a dose-dependent increase in Cten, ROCK1, Src, Snail, and N-cadherin expression, whereas the protein expression level of E-cadherin was decreased following stimulation with TGF-β1 (Figure 1 A). The optimum concentration of TGF-β1-human recombinant protein (20 ng/ml) was selected for subsequent TGF-β1 stimulation experiments.

**Figure 1:**
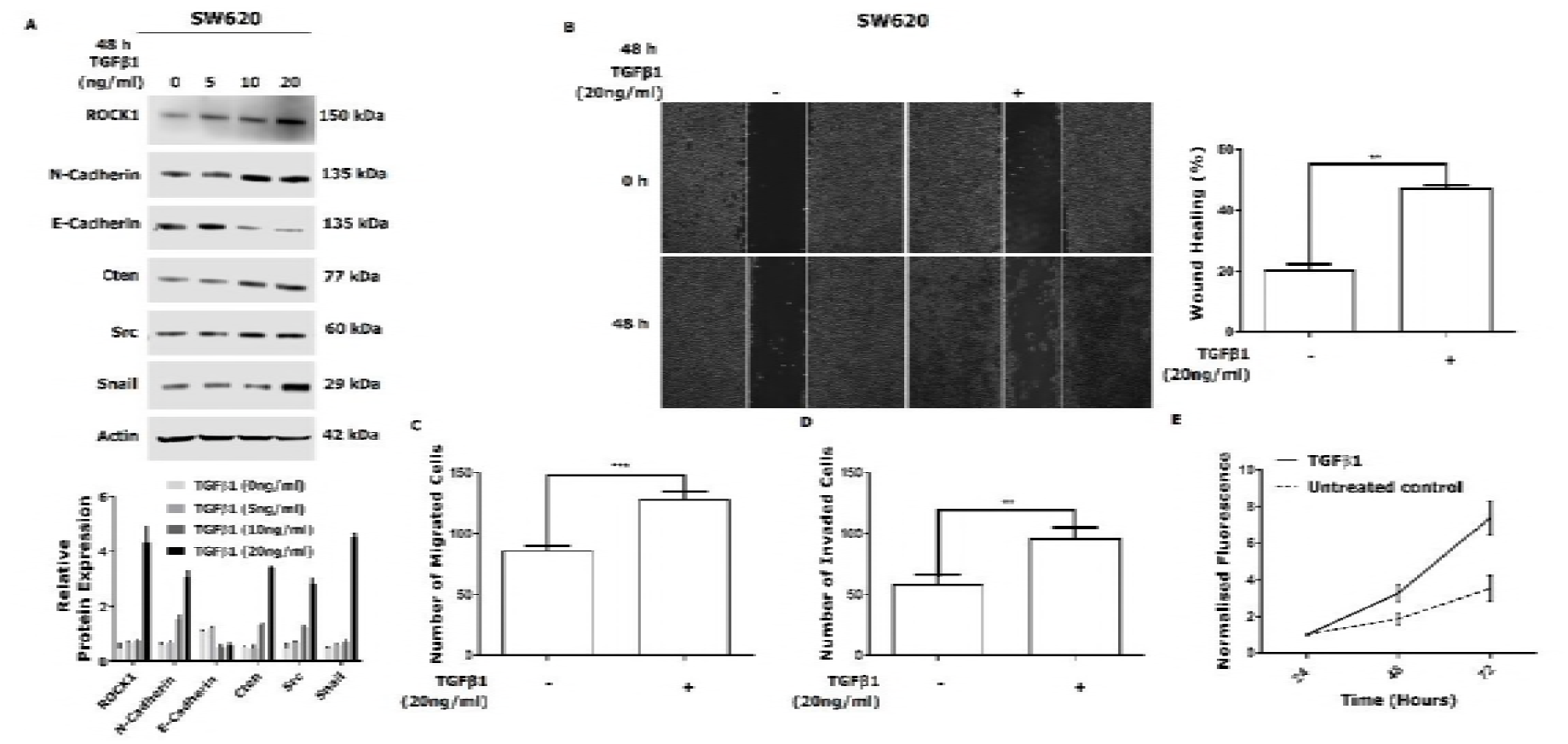
TGFβ1 increases Cten protein expression in a dose-dependent manner. A) SW620 cells were stimulated with TGFβ1-human recombinant protein (R&D systems), (0-20 ng/ml) for 48 hours and the changes in Cten, ROCK1, N-cadherin, E-cadherin, Src, and Snail protein expression were determined by western blot (upper panel). Graph on the lower panel represents the densitometry values calculated for each protein band normalised to actin. Stimulation of SW620 cells with TGFβ1 treatment (20 ng/ml for 48 hours) induced closure of wound compared to untreated control (P = 0.0024). C) Stimulation of TGFβ1 was associated with an increase in cell migration compared to untreated SW620 cells control (P = 0.0005). D) Treatment of SW620 cells with TGF-β1-human recombinant protein enhanced cell invasion compared to untreated control (P = 0.0055). E) Stimulation of TGFβ1 in SW620 cells was associated with an increase in cell proliferation compared to untreated control (P ≤ 0.0001). Results are representative of at least 3 experimental replicates.**=P<0.01

The relationship between TGF-β1 and Cten was further investigated in an additional cell line, HCT116. In agreement with the findings in SW620 cells, stimulation of TGF-β1 was associated with an increase in the protein expression levels of Cten, ROCK1, Src, Snail, and N-cadherin, whereas E-cadherin expression was inhibited (Figure 2 A). The ability of TGF-β1 to alter cell motility and viability was then investigated using the Transwell migration assay, wound healing assay, Transwell Matrigel invasion assay, and cell viability assay. In both SW620 and HCT116 cell lines, cell migration, invasion, and proliferation were increased when cells were treated with TGF-β1-human recombinant protein compared to untreated control (Figure 1B,C,D,E), (Figure 2B,C,D,E).

**Figure 2:**
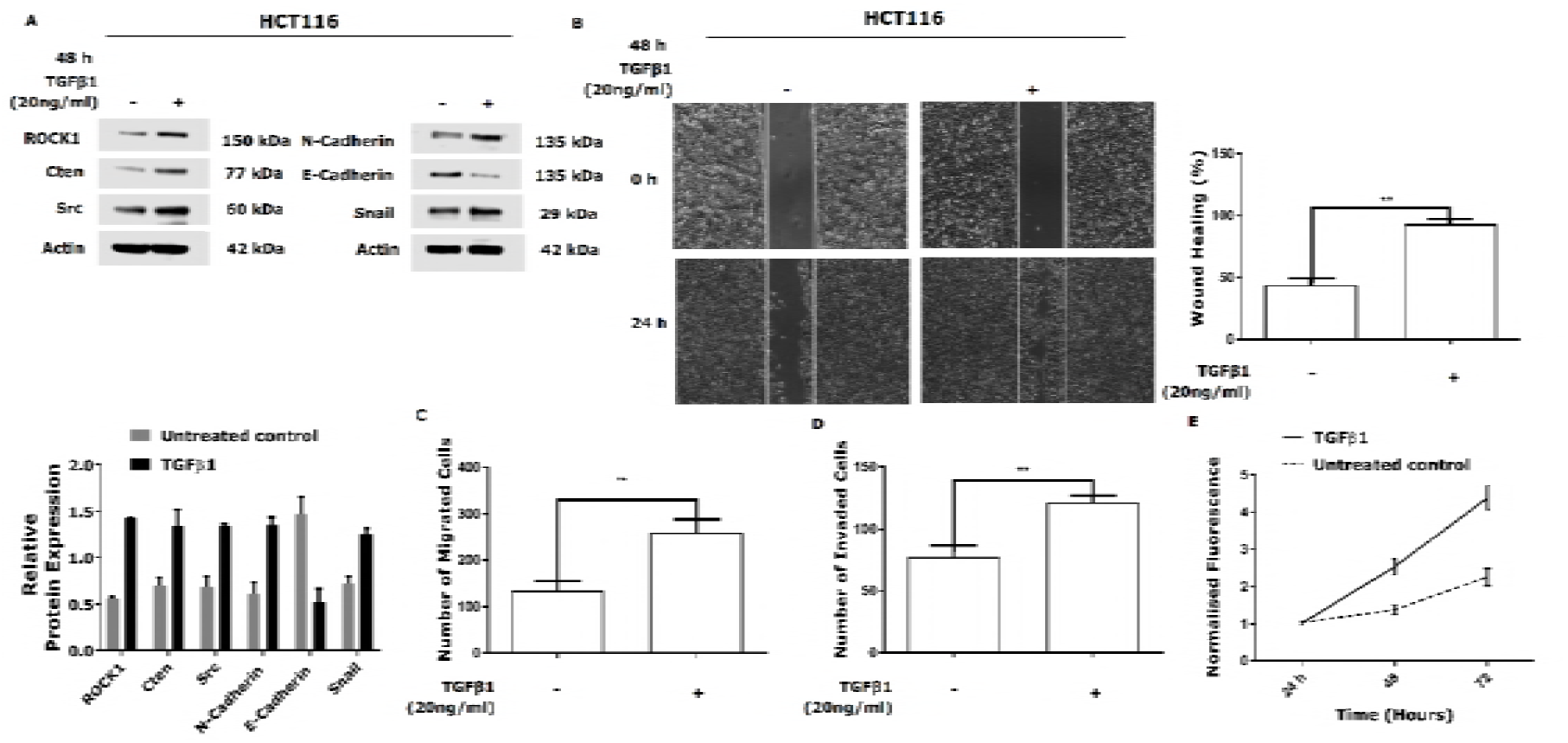
TGFβ1 stimulation increases Cten protein expression in HCT116 cells. A) HCT116 cells were pre-treated with TGF-β1-human recombinant protein (20 ng/ml for 48 hours) and the changes in Cten, ROCK1, N-cadherin, E-cadherin, Src, and Snail protein expression were determined by western blot (upper panel). Graph on the lower panel represents the densitometry values calculated for each protein band normalised to actin. B) Treatment of HCT116 cells with TGFβ1 TGF-β1-human recombinant protein (20 ng/ml for 48 hours) induced wound closure compared to untreated control (P = 0.0084). C) Stimulation of TGFβ1 was associated with an increase in cell migration compared to untreated HCT116 cells control (P = 0.0032). D) Stimulation of SW620 cells with TGFβ1 treatment increased cell invasion compared to untreated control (P = 0.0022). E) Stimulation of TGFβ1 in HCT116 cells resulted in an increase in cell proliferation compared to untreated control (P ≤ 0.0001). Results are representative of at least 3 experimental replicates.**=P<0.01

The effect of TGF-β1 on Cten and its downstream targets was again investigated in SW620 cell but using an alternative methodology. Assuming that there was some endogenous production of TGF-β1 by the cell lines, this was transiently knocked down by siRNA and changes in protein expression level of Cten, ROCK1, Src, Snail, E-cadherin and N-cadherin were evaluated by western blot. Confirming the TGF-β1 stimulation results, knockdown of TGF-β1 resulted in a reduction of Cten, ROCK1, Src, Snail, N-cadherin protein expression levels. Additionally, TGF-β1 knock down was associated with an increase in the expression level of E-cadherin compared to luciferase targeting siRNA control (Figure 3 A). The effect of TGF-β1 knockdown on cell functions was also tested in this study (Figure 3B,C,D,E). Knockdown of TGF-β1 was associated with a significant decrease in cell migration, invasion and proliferation compared to the luciferase control (Figure 3). Collectively, these findings suggest that TGF-β1 may promote both cell motility and proliferation through the upregulation of EMT processes in CRC cells.

**Figure 3:**
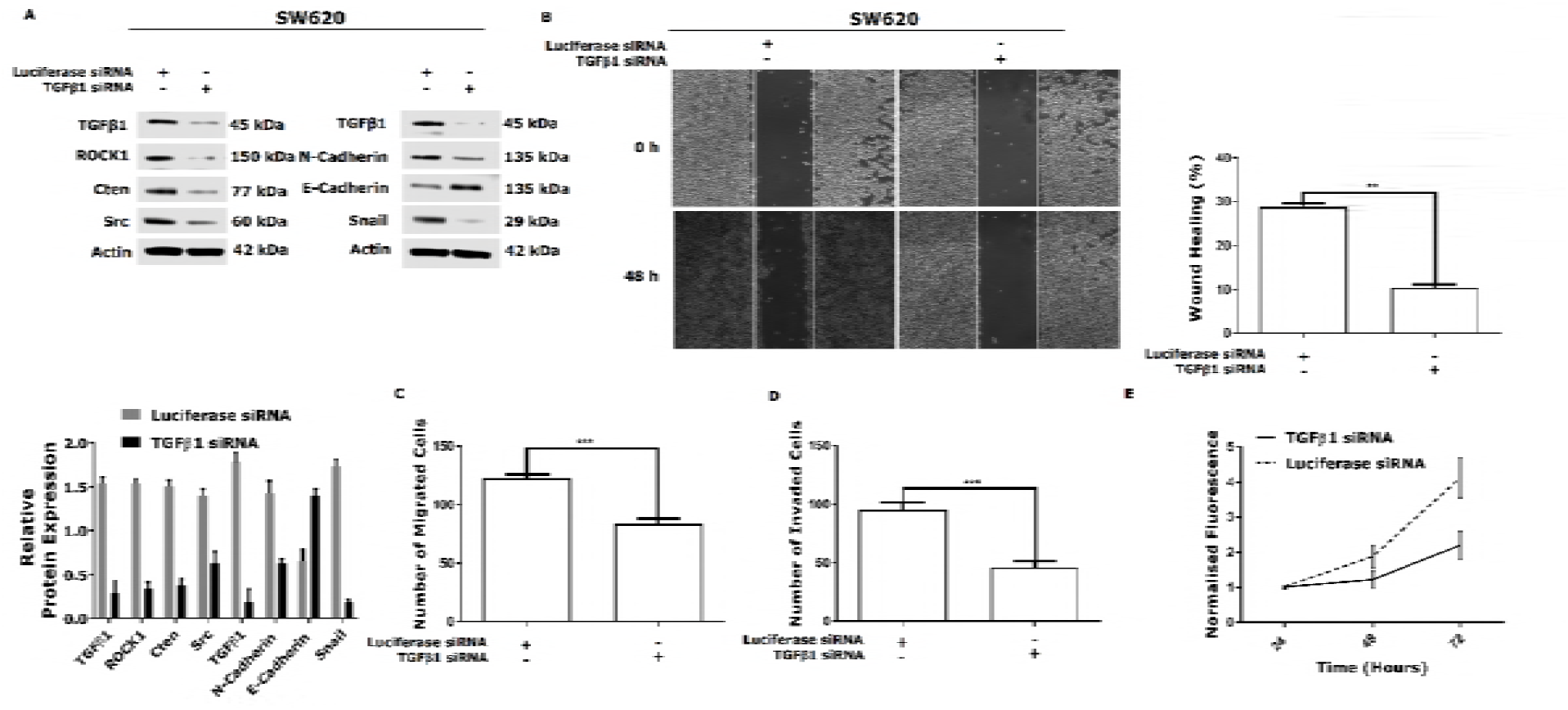
TGFβ1 knockdown decreases Cten protein expression in SW620 cells. A) SW620 cells were transfected with TGFβ1 targeting siRNA duplexes (200 nM/ml for 48 hours) and the changes in Cten, ROCK1, N-cadherin, E-cadherin, Src, and Snail protein expression were determined by western blot (upper panel). Graph on the lower panel represents the densitometry values calculated for each protein band normalised to actin. B) Knockdown of TGFβ1 in SW620 decreased wound closure compared to luciferase targeting siRNA control (P = 0.0016). C) Knockdown of TGFβ1 was associated with a decrease in cell migration compared to luciferase transfected HCT116 cells control (P = 0.0003). D) Knockdown of TGFβ1 in SW620 cells decreased cell invasion compared to luciferase siRNA control (P = 0.0005). E) Knockdown of TGFβ1 in SW620 cells resulted in a reduction in cell proliferation compared to luciferase siRNA control (P ≤ 0.0001). Results are representative of at least 3 experimental replicates. **=P<0.01, ***=P<0.001

### TGFβ1 Induces Nuclear Localisation of Cten/Src/ROCK1/Snail

Cten is normally located at focal adhesions but we and others have shown that there is also nuclear localisation (although the significance of this is uncertain). Since TGF-β1 signalling may be mediated through Cten, it was of interest whether TGF-β1 is capable of inducing Cten protein translocation to the nucleus. HCT116 cells were stimulated with TGF-β1 and the expression of Cten and its downstream targets in nucleus were observed by immunofluorescences staining (Figure 5A). The results showed that stimulation of HCT116 cell with TGF-β1 for 48 h induced Cten, Src, ROCK, and Snail protein expression as well as translocation to the nucleus. In contrast, in untreated HCT116 cells, there was no Cten or its downstream target proteins detected in the nucleus (Figure 5A). To validate these data, TGF-β1 was used to stimulate the SW620 cell line and both immunofluorescence staining and nuclear and cytoplasmic extraction were performed to directly determine the subcellular localisation of Cten and its downstream targets: Src, ROCK1, and Snail proteins. Subcellular fractionation experiments showed an increase in the nuclear fraction of Cten, Src, ROCK1, or Snail proteins following treatment with TGF-β1 (Figure 4). Immunofluorescence imagining showed that, as with HCT116, stimulation of SW620 with TGF-β1 treatment increased the expression and nuclear translocation of Cten, ROCK1, Src, and Snail compared to the untreated cell control (Figure 5B).

**Figure 4:**
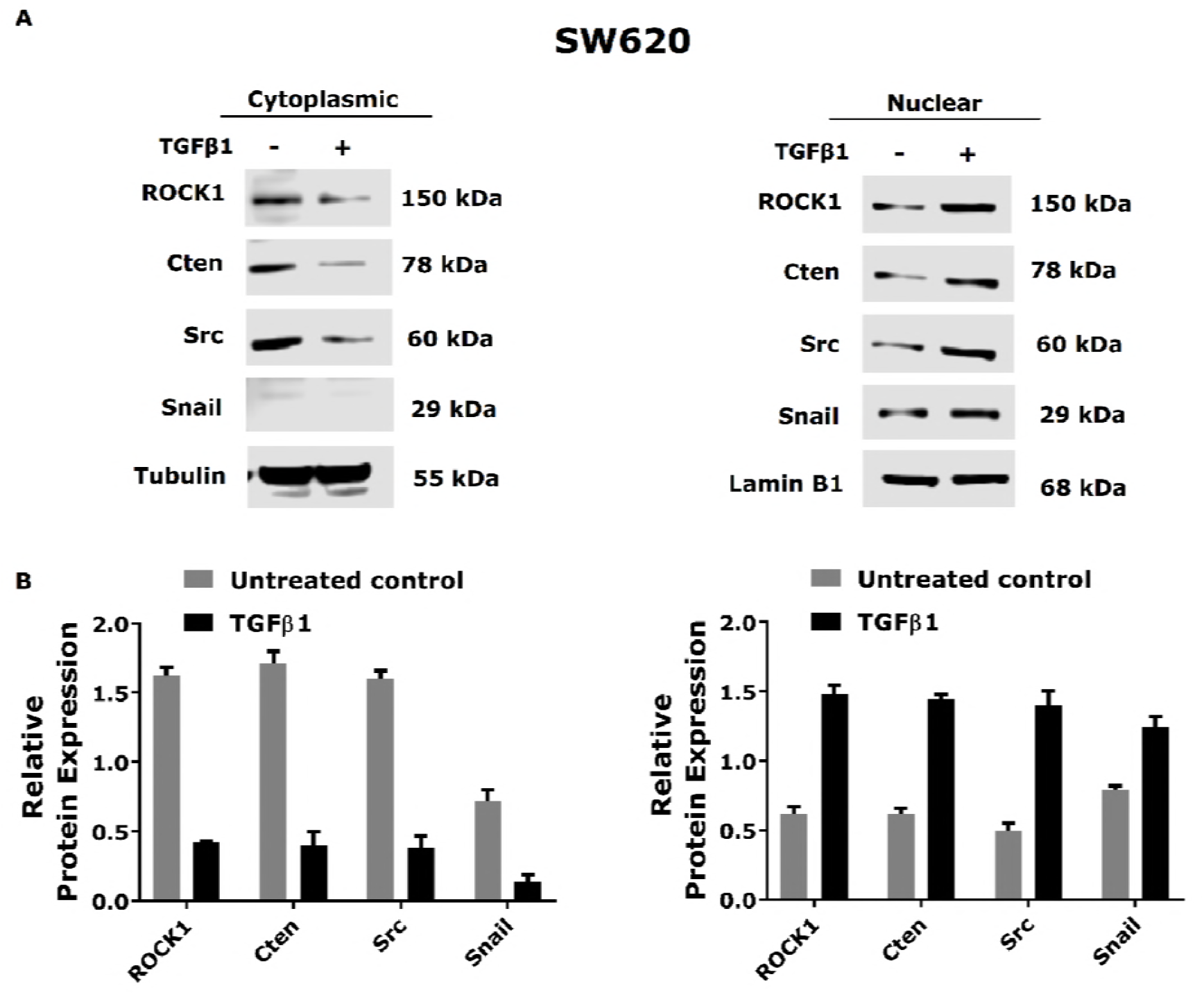
TGFβ1 induces nuclear translocation of Cten and its downstream targets in SW620 cells. A) Subcellular fractionation extraction was performed following the treatment of SW620 cell with or without 20 ng/ml of TGF-β1-human recombinant protein for 48 hours and the lysates were assessed by western blot. Lamin B1 and tubulin were used as loading control for cytoplasmic and nuclear fractions. B) Quantitative determination of the relative expression of ROCK1, Cten, Src, and Snail protein fractions following treatment with TGFβ1.

**Figure 5:**
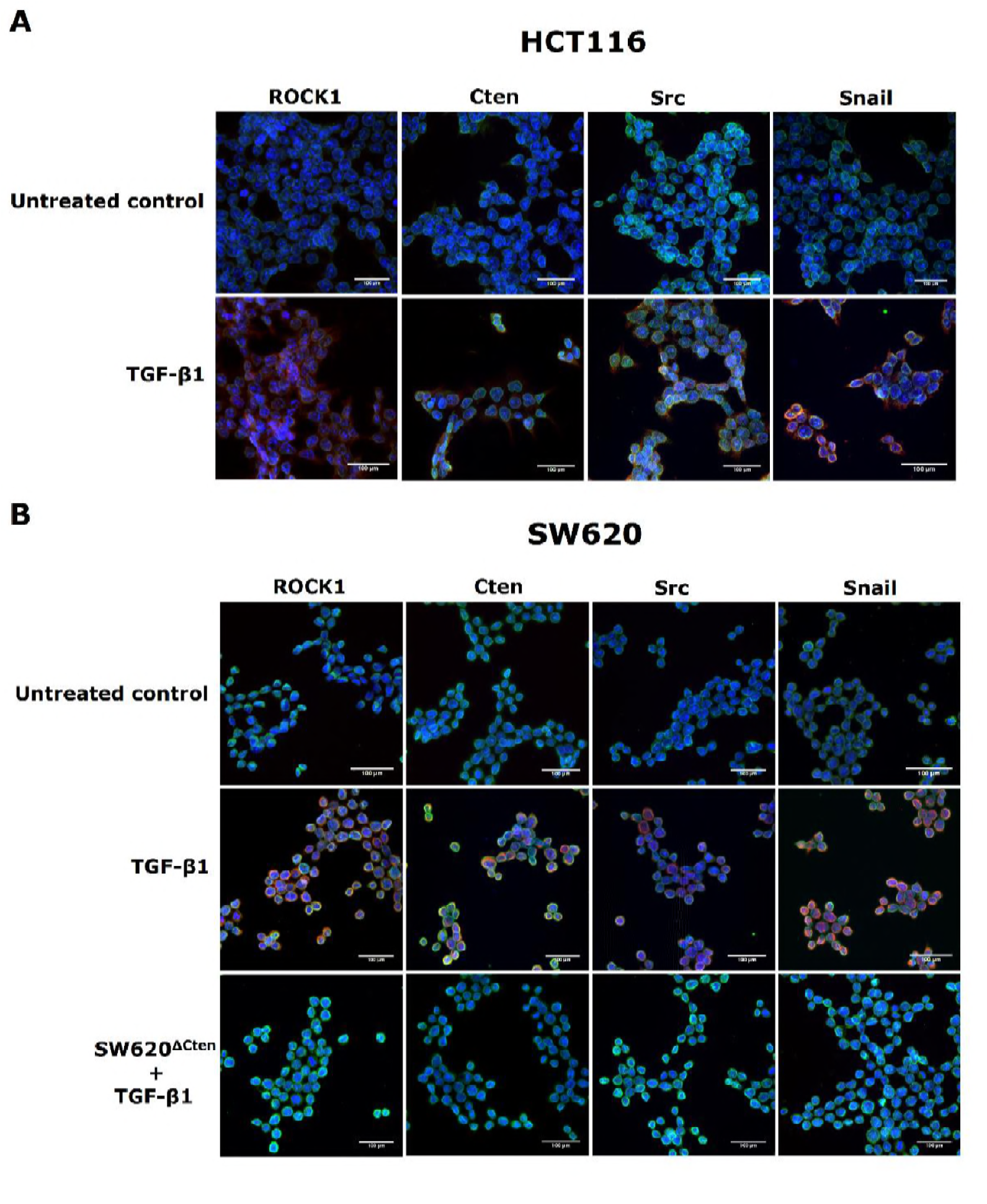
TGFβ1 induces nuclear localisation of Cten/Src/ROCK1/Snail. A and B) Stimulation of TGFβ1 increases nuclear localisation and expression of Cten and its downstream targets in HCT116 and SW620 cells but this was lost when Cten was knocked out in SW620^ΔCten^ cells. The subcellular localisation of ROCK1, Src, Cten, and Snail was examined by confocal microscopic images of DAPI (blue), nuclear envelope marker Lamin B (green) and ROCK1, Src, Cten or Snail protein (17) in untreated control and treated cells with TGFβ1 stimulation (20ng/ml) for 48 hours. (scale bar100 μm).

To investigate whether Cten was essential for TGF-β1 induced the nuclear translocation of Src, ROCK and Snail proteins through Cten, the cell line SW620^ΔCten^, was stimulated with TGF-β1 and the nuclear translocation of the protein was determined by immunofluorescence. The results revealed that, in the absence of Cten, the nuclear translocation of Src, ROCK and Snail following TGF-β1 stimulation (Figure 5 B), does not occur. Taken together, these data suggest that nuclear translocation of ROCK1, Src, and Snail protein is probably mediated by TGF-β1 via the upregulation of the Cten signalling pathway

### Cten deletion selectively abrogates TGF-β1 Induced Cell Migration and Invasion

Our data have shown that TGFβ1 is a positive regulator of Cten expression and a number of markers of EMT. We also showed that TGFβ1 signalling induces cell motility (both migration and invasion). Since these Cten induces EMT and cell motility, it is reasonable to hypothesise that Cten in a direct mediator of TGFβ1 activity. We have previously described the creation of SW620^ΔCten^ (16). This is derivative of SW620 in which Cten has been deleted using CRISPR/Cas9 technology. In order to test our hypothesis, SW620^ΔCten^ was stimulated with TGFβ1and Western blotting was performed. The results revealed that stimulation with with TGF-β1 was associated with a small increase in N-cadherin expression but ROCK1, Src, Snail, and E-cadherin protein expression level remained unchanged compared to the sham-treated cells control (Figure 6 A). This implies that Cten is may be responsible for TGF-β1 induced EMT.

**Figure 6:**
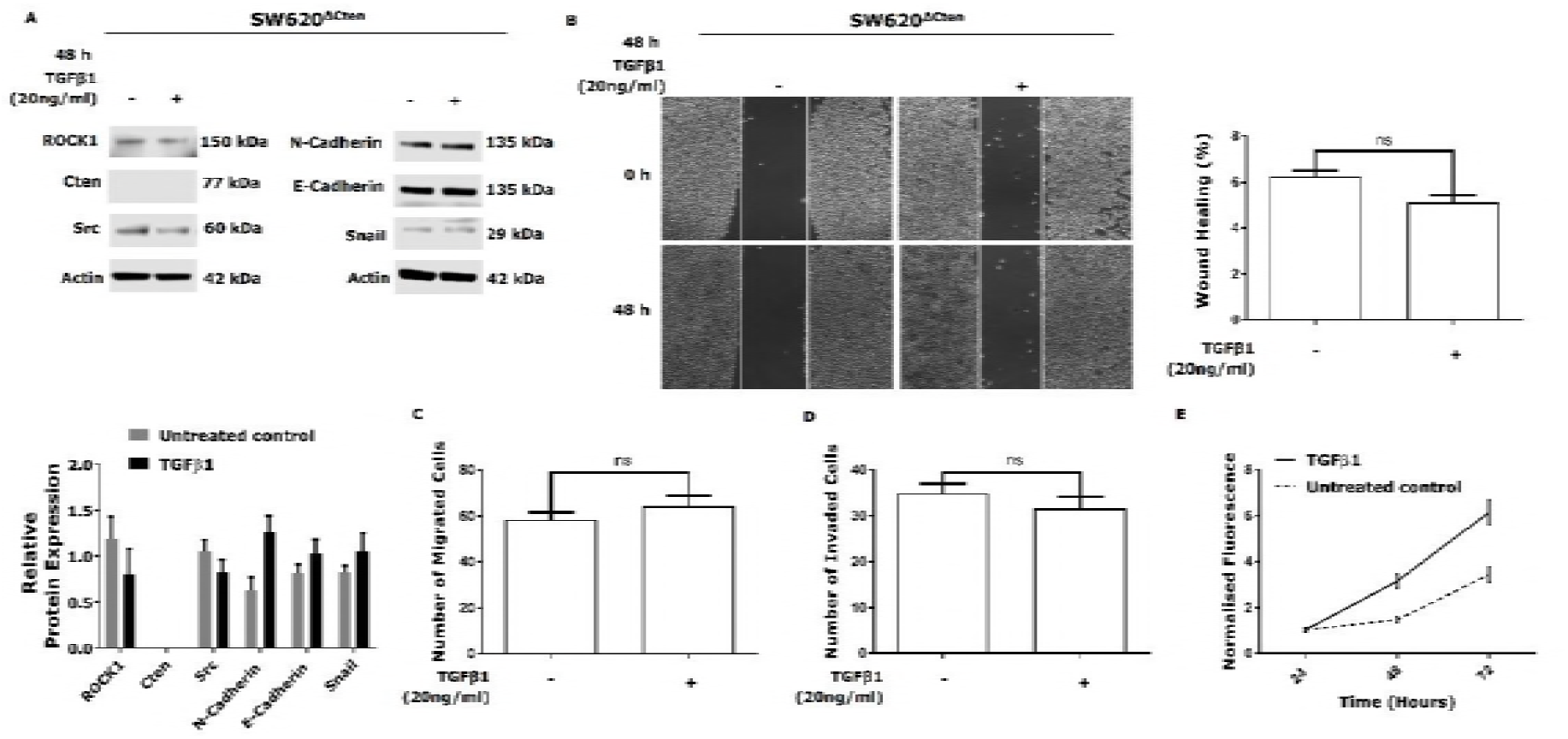
TGFβ1 signals through Cten to regulates EMT and promotes cell migration and invasion. A) Stimulation of SW620^ΔCten^ cells with TGFβ1 treatment (20 ng/ml for 48 hours) was associated with a small increase in N-cadherin expression and ROCK1, Src, Snail, and E-cadherin protein expression level remained unchanged from the untreated cells control (upper panel). Graph on the lower panel represents the densitometry values calculated for each protein band normalised to actin. B) Wound healing assay showed no significant differences between TGFβ1 stimulation and untreated SW620^ΔCten^ cells control (P = 0.0585). C) Stimulation of TGFβ1 in SW620^ΔCten^ cells did not cause a significant increase in cell migration compared to untreated control (P = 0.1561). D) Treatment of SW620^ΔCten^ cells with TGF-β1-human recombinant protein did not enhance cell invasion compared to untreated control (P = 0.1469). E) Stimulation of TGFβ1 in SW620^ΔCten^ cells was associated with an increase in cell proliferation compared to untreated control (P ≤ 0.0001). Results are representative of at least 3 experimental replicates. **=P<0.01

To determine if Cten was functionally relevant to TGF-β1 mediated activity, SW620^ΔCten^was stimulated TGF-β1 and a assays for Transwell migration, wound healing, Matrigel invasion, and cell viability were performed. (Figure 4B,C,D,)The deletion of Cten in SW620^ΔCten^ cells resulted in an abrogation of the ability of TGF-β1 to induce cell migration or invasion whilst the ability to induce cell proliferation was retained. Thus Cten appears not to be involved in TGF-β1 induced cell proliferation, but it may be a signalling intermediate in the TGF-β1/EMT pathway regulating cell migration and invasion in CRC cells.

## DISCUSSION

EMT is a critical process occurring during tumour metastasis. During the EMT process, epithelial cells show loss of cell to cell adhesion by E-cadherin downregulation at adherens junctions, cytoskeleton reorganisation via switching from keratin to vimentin intermediate filaments, loss of apical-basal polarity, acquisition of mesenchymal cell phenotype and increased cell invasion and migration (18). Cten has been shown to act as oncogene in most tumour types and involved in regulation of EMT processes, however, the mechanisms that upregulate the expression of Cten induced EMT have not been elucidated (16, 19–21). Previous research from our laboratory suggested that Cten expression is regulated by EGFR/KRAS signalling (5) and the study by Katz et al (3) showing the role of Her2 in up-regulating Cten would seem to validate this. We and others further have also found that Cten could be under the regulation of several cytokines such as IL6/Stat3 and growth factors (6, 22). The present study directly shows, for the first time an essential role for Cten in TGF-β1-induced EMT and cell motility in CRC cells and that this may be through upregulation of the Src/ROCK1/Snail signalling pathway.

TGF-β1 reputedly plays a key role in promoting EMT initiation and tumour metastasis. In the latest classification of CRCs (25), the most aggressive class (CMS4) is characterised by TGF-β activation. It has been documented that TGF-β1 induces the expression of several transcription factors including Twist, Zeb, Slug, and Snail (23). Here, using different methods of modulating TGF-β1 activity, we have shown that TGF-β1 is a positive regulator of Cten expression. We have also shown that TGF-β1 is a positive regulator of Src, ROCK, and Snail expression. Since these are putative targets of Cten, it begs the question whether, in this case, they are directly up-regulated by TGF-β1 signalling or whether they are up-regulated by Cten as a secondary event. Using the SW620^ΔCten^ cell line (in which Cten is deleted) we were able to conclusively show that up-regulation of these molecules is Cten-dependent. We observed, by both immunofluorescence and cellular fractionation, that TGF-β1 signalling induced nuclear localisation of Cten as well as Src, ROCK, and Snail. We were similarly able to show that nuclear localisation of Src, ROCK, and Snail was Cten-dependent. The mechanisms by which these proteins are up-regulated and by which their cellular location is controlled are uncertain although we have previously shown that Cten causes post-translational stabilisation of Snail (16). The link between TGF-β1 and Cten may be through integrins since TGF-β1 induces integrin activation and Cten is found in compelxt with the cytoplasmic tail of integrins. However there are also several other downstream targets of TGF-β1 mediated EMT which may possibly be involved in regulation of Cten and further investigation of markers/signalling pathways such as Ras/MAPK (10), RhoA (11), and Jagged 1/Notch (12) is warranted.

Our data suggest a novel downstream signalling pathway for TGF-β1 which is mediated through Cten. In order to ascertain whether this represented a functionally relevant pathway, we performed assays to assess motility (both by invasion and migration assays) and cell proliferation. In our models, TGF-β1 signalling was shown to increase cell proliferation, migration (both transwell migration and wound healing) and cell invasion. These data are in accordance with previous studies that TGF-β1 induced EMT can promote cell motility and proliferation in a vast range of different tumour cells such as breast cancer and oral squamous cell carcinoma (24, 26). Intuitively one would think that the induction of cell motility was mediated through the Cten pathway. This was confirmed by our observation that there was a selective loss of the effect of TGF-β1 treatment on both cell migration and invasion in the SW620^ΔCten^ cell line while the effect of TGF-β1 treatment on cell proliferation was unaffected. This is completely in line with our previous data showing that Cten regulates cell motility and does not affect cell proliferation (19).

It is not unexpected that TGF-β1will activate different signalling pathways to regulate different cellular functions. Our data show that the Cten pathway is an important factor in the TGF-β1 regulation of cell motility and they may explain other observations that have been made. Thus previous published data from our group have shown that Cten induces EMT and promotes cell motility through FAK, ILK and Snail signalling (16, 21, 27). Others have shown that FAK and/or ILK are required for TGF-β1 induced EMT and to promote cell motility (13, 28). Snail also has been shown to act as a mediator of TGF-β1 induced EMT(29). We would now hypothesise that Cten is one of the missing links which would explain FAK/ILK/Snail dependence of TGF-β1 induced cell motility.

In addition to Cten, there are other downstream pathways involved in TGF-β1 mediated cell migration, including JAK/STAT3 (30), PI3K-Akt (31), and Reelin (32), so it would be of interest to determine whether Cten acts in parallel or synergistically with these pathways in future studies

In summary, the data presented have indicated that TGFβ1 and Cten signalling may cooperate in promoting EMT and cell motility. Regulation of downstream markers of EMT such as Src/ROCK1/Snail by TGFβ1 is dependent on Cten. These processes are relevant to the development of metastasis and our data open up the possibility of targeting Cten in CRC.

## Funding

This work was supported by the Saudi Arabian Cultural Bureau in Britain (S3555).

## Conflict of interest

The authors declare that they have no conflicts of interest with the contents of this article

